# Perception of rhythmic speech is modulated by focal bilateral tACS

**DOI:** 10.1101/647982

**Authors:** Benedikt Zoefel, Isobella Allard, Megha Anil, Matthew H Davis

## Abstract

Several recent studies have used transcranial alternating stimulation (tACS) to demonstrate a causal role of neural oscillatory activity in speech processing. In particular, it has been shown that the ability to understand speech in a multi-speaker scenario or background noise depends on the timing of speech presentation relative to simultaneously applied tACS. However, it is possible that tACS did not change actual speech perception but rather auditory stream segregation. In this study, we tested whether the phase relation between tACS and the rhythm of degraded words, presented in silence, modulates word report accuracy. We found strong evidence for a tACS-induced modulation of speech perception, but only if the stimulation was applied bilaterally using ring electrodes (not for unilateral left hemisphere stimulation with square electrodes). These results were only obtained when data was analyzed using a statistical approach that was identified as optimal in a previous simulation study. The effect was driven by a phasic disruption of word report scores. Our results suggest a causal role of neural entrainment for speech perception and emphasize the importance of optimizing stimulation protocols and statistical approaches for brain stimulation research.

## Introduction

Neural oscillations align to rhythmic input, a mechanism termed neural entrainment (Lakatos, Karmos, Mehta, Ulbert, & Schroeder, 2008). Neural entrainment is proposed to be a crucial mechanism underlying speech processing (Ding & Simon, 2014; Peelle & Davis, 2012; Zoefel & VanRullen, 2015b). Using transcranial alternating current stimulation (tACS), we can selectively perturb neural oscillations (Herrmann, Rach, Neuling, & Strüber, 2013) and observe the consequences for perception and comprehension. Previous studies have used tACS to suggest a causal role of neural entrainment for speech processing. Combining left-lateralised tACS over auditory brain regions with functional magnetic resonance imaging (fMRI), Zoefel, Archer-Boyd, & Davis (2018) showed that the phase relation between tACS and rhythmic speech impacts the blood-oxygen level dependent (BOLD) response to intelligible but not to matched, unintelligible speech. Two other tACS studies found that bilateral neural modulation also impacts on word report for spoken sentences; Wilsch, Neuling, Obleser, & Herrmann (2018) and Riecke, Formisano, Sorger, Başkent, & Gaudrain (2018) both found that word report accuracy (a measure of speech perception success) depends on the time delay between envelope-shaped tACS stimulation and speech signals. While these findings are striking and provide compelling evidence that tACS modulates neural activity and speech perception in parallel, they still fall short of demonstrating a causal role of oscillatory neural entrainment on speech perception. Primarily, this is because the target sentences in the experiments reported by Wilsch et al. (2018) and Riecke et al. (2018) were masked by background noise or other speech. The success of speech perception in noise depends on listener’s ability to segregate speech and background sounds (i.e. auditory scene analysis; Pressnitzer, Suied, & Shamma, 2011), as well as on speech processing per-se. It might be that tACS improved auditory scene analysis (in line with a previous study by Riecke, Sack, & Schroeder, 2015) instead of modulating speech perception directly.

In this study, we therefore tested whether we can use tACS to modulate the success of speech perception rather than auditory scene analysis. Similar to some of the previous studies cited above (Experiment 1 in Riecke et al., 2018; Zoefel, Archer-Boyd, et al., 2018), we systematically varied the phase relation between tACS and rhythmic speech (Fig. 1A). Instead of presenting two competing speakers as in Riecke et al. (2018), we used rhythmic sequences of isolated words whose intelligibility was manipulated using noise-vocoding (Shannon, Zeng, Kamath, Wygonski, & Ekelid, 1995) as in the tACS-fMRI study reported by Zoefel, Archer-Boyd, et al. (2018). By using intrinsic acoustic degradation (cf. Mattys, Davis, Bradlow, & Scott, 2012) of speech presented in a silent background, we can ensure that effects of tACS are due to modulation of speech perception per-se, and not due to effects on other processes involved in auditory scene analysis, or selective attention.

**Figure 1.**
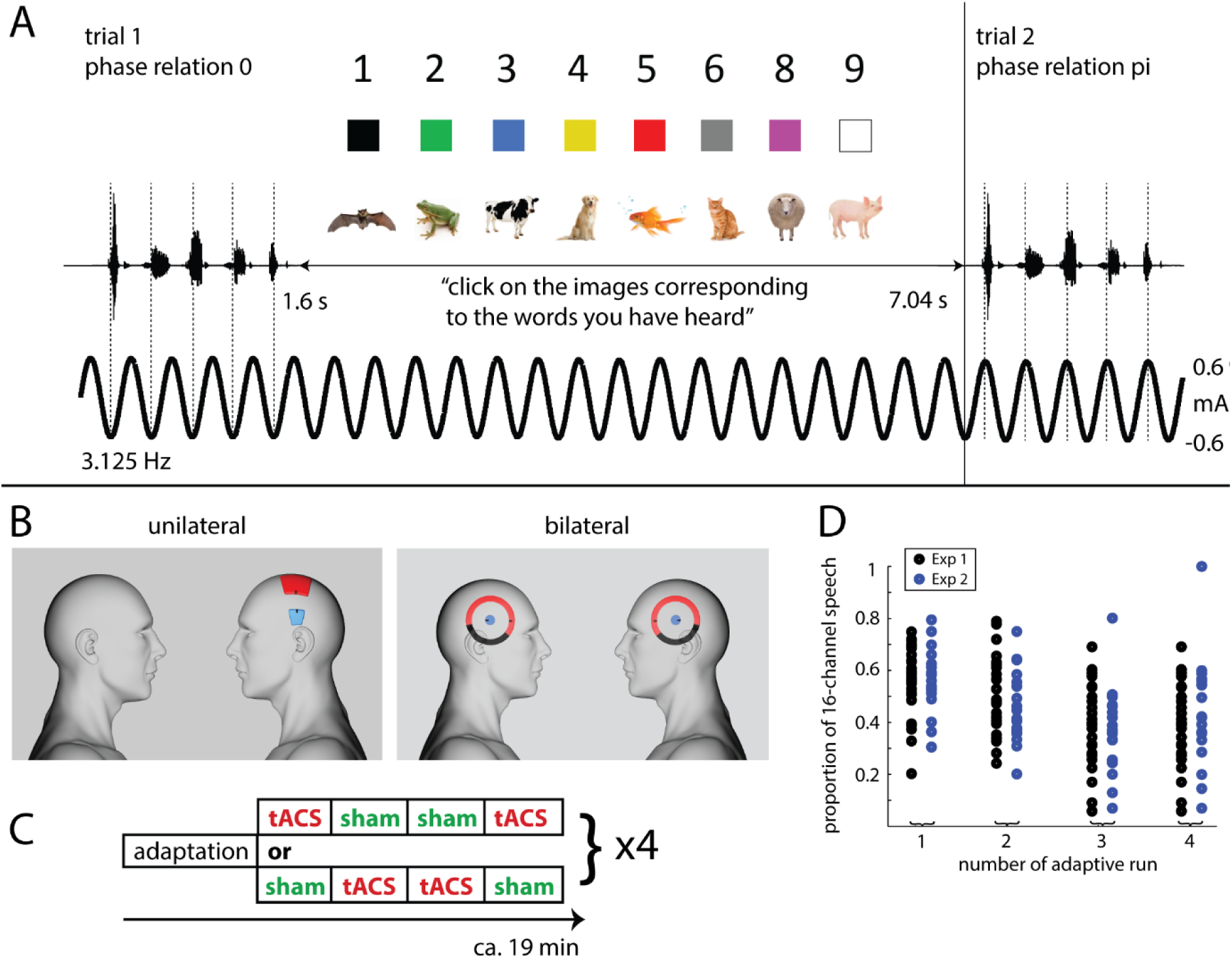
Experimental paradigm. A. Rhythmic noise-vocoded speech sounds, consisting of five one-syllable words, were presented so that the p-center of all syllables (dashed lines) fell at one of eight different phases of simultaneously applied tACS. After each trial, participants selected images that correspond to the second, third and fourth word they thought to have heard, with eight possible options for each word. B. Electrode configuration in Experiment 1 (unilateral) and 2 (bilateral). For both configurations, connector positions are shown in black. For the bilateral configurations, parts of the outer electrodes covered by isolating tape are colored black. C. Blocked design used in both experiments. Participants completed 4 x 5 runs, with two possible run orders as shown here and described in detail in Section Experimental Design. D. Proportion of 16-channel vocoded speech used to construct vocoded stimuli (for which 16- and 1-channel speech were mixed; see Section Stimuli), separately for each participant and the four adaptive runs in the two experiments.

Previous tACS studies (e.g., Riecke et al., 2018; Wilsch et al., 2018) focused on the demonstration of a phasic modulation of speech perception outcomes (and therefore a functional consequence of oscillatory brain stimulation). However, for real-world applications of tACS, the most important outcome is clearly to enhance speech perception relative to an unstimulated control condition (Peelle, 2018; Riecke & Zoefel, 2018). It is equally likely that the reliable effect of tACS is to disrupt word report relative to an unstimulated condition, but previous work remained inconclusive in this respect (summarized in the Discussion). By using the Log Odd’s Ratio (LOR) to quantify the word report difference between stimulation and sham conditions, and by phase-aligning to both maximum and minimum LOR, we therefore separately assessed whether tACS enhances and/or disrupts word report accuracy relative to sham.

We also compared two different stimulation protocols in their efficacy of modulating speech perception (Fig. 1B): The first (Experiment 1), using standard square electrodes and unilateral stimulation over left Superior Temporal Gyrus (STG), is identical to that which was combined with fMRI by Zoefel, Archer-Boyd, et al. (2018). The second (Experiment 2) consisted of bilateral stimulation over STG, using ring electrodes, shown to improve the focality of the stimulation (Datta et al., 2009; Saturnino, Antunes, & Thielscher, 2015; see also Heise et al., 2016).

## Methods

### Participants

Forty-seven volunteers were tested after giving informed consent under a process approved by the Cambridge Psychology Research Ethics Committee. Twenty-seven volunteers (15 females; mean ± SD, 31 ± 7 years) participated in Experiment 1, twenty volunteers participated in Experiment 2. One volunteer did not finish Experiment 2 as they did not feel comfortable with the stimulation, leaving nineteen volunteers (8 females; 21 ± 2 years). All volunteers were native English speakers and had no history of hearing impairment, neurological disease, or any other exclusion criteria for tACS based on self-report.

### Stimuli

We used rhythmic noise-vocoded speech; the same stimulus as in a previous tACS-fMRI study (Zoefel, Archer-Boyd, et al., 2018). The original speech consisted of sequences of 5 one-syllable words, spoken in time with a metronome beat at 2 Hz by a male native speaker of Southern-British English (MHD). This approach resulted in an alignment of the speech’s “perceptual centers”, or “p-centers” (Morton, Marcus, & Frankish, 1976), with the metronome beat, and consequently perfectly rhythmic speech sequences. We assumed that the rhythmicity of the stimulus would promote neural entrainment; it also enabled us to define phase relations between speech and tACS in a straightforward manner (see below).

The original speech was time-compressed to 3.125 Hz using the PSOLA algorithm included in Praat software (version 6.12, from http://www.fon.hum.uva.nl/praat/download_win.html).

Individual words were extracted and combined into sentences of five words, in the following order: “pick” <number> <colour> <animal> “up”, where <number> could be any number between one and nine (excluding the bi-syllable seven);<colour> could be any of the following: “black”, “green”, “blue”, gold”, “red”, “grey”, “pink”, “white”; and <animal> could be any of the following: “bat”, “frog”, “cow”, “dog”, “fish”, “cat”, “sheep”, “pig”.

These time-compressed five-word sentences were then manipulated using noise-vocoding (Shannon et al., 1995), a technique which can be used to alter the intelligibility of speech without strongly affecting the broadband envelope which is hypothesized to be critical for neural entrainment (Ghitza, Giraud, & Poeppel, 2013). The speech signal was first filtered into 16 logarithmically spaced frequency bands between 70 and 5000 Hz, and the amplitude envelopes (Fig. 2A, top) were extracted for each band (half-wave rectified, low-pass filtered below 30 Hz). The envelope for each of those frequency bands, *env*(*b*), was then mixed with the broadband envelope (Fig. 2A, bottom) of the same speech signal, *env*(*broadband*), in proportion *p,* to yield an envelope for each frequency band, *env_final_*(*b*). These constructed envelopes (Fig. 1B) were then used to construct the final speech stimuli.

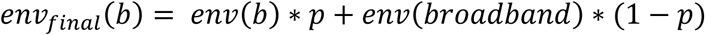

**Figure 2.**
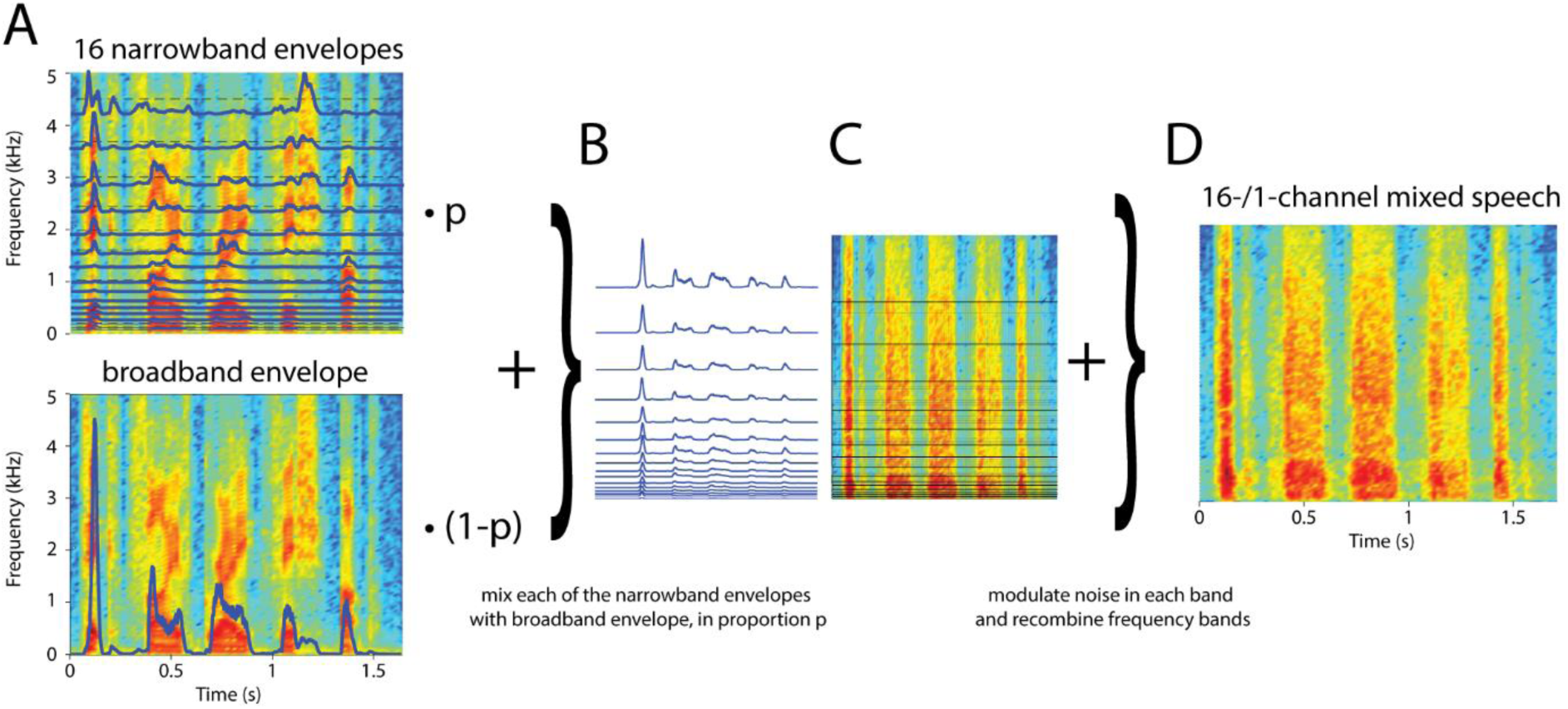
Stimulus construction. A. For each of the clear speech sentences (spectrogram for one example sentence is color-coded and shown in both panels), amplitude envelopes (blue lines) were extracted for 16 frequency bands (top) as well as the broadband signal (bottom). B. Each of the 16 narrowband envelopes was mixed with the broadband envelope in proportion p (0.5 for the example shown in B-D). C. Each of the resulting envelopes (shown in B) was used to modulate noise in the respective frequency band. D. The resulting signals were re-combined to yield a 16-/1-channel vocoded speech mix. For this form of vocoded speech, high values of the mixing proportion (p = 1) results in 16-channel vocoded speech which is highly intelligible, low mixing proportions (p = 0) results in 1-channel vocoded speech which is entirely unintelligible. Intermediate proportions result in intermediate intelligibility.

If *p* is 0, *env_final_* is identical for each frequency band, and identical to the broadband envelope. This creates an unintelligible, 1-channel stimulus, similar to unintelligible speech conditions used in previous brain imaging (Peelle, Gross, & Davis, 2013), tACS (Zoefel, Archer-Boyd, et al, 2018), and behavioural studies (Sohoglu, Peelle, Carlyon, & Davis, 2014). If *p* is 1, *env_final_*(*b*) is identical to *env*(*b*). For 16-channel vocoded speech, this leads to nearly full intelligibility for spoken words in typical sentences, as used in previous studies (ibid). For intermediate values (between 0 and 1), our manipulation has the effect of gradually increasing the degree of co-modulation (i.e. correlation between envelopes) of the different frequency bands of vocoded speech, ranging from perfectly co-modulated (0% morphing, corresponding to 1-channel vocoded speech) to a co-modulation equivalent to that seen for “standard” 16-channel vocoded speech. The constructed envelopes (*env_final_*) were applied to broad-band noise (Fig. 2C), filtered using the same logarithmically spaced filters as used in analysis, and the output signals were re-combined to yield noise-vocoded speech stimuli (Fig. 2D). Importantly, the intelligibility of these stimuli varies with *p* (the speech sounds like noise and is completely unintelligible if *p* is 0, and sounds like clearly intelligible speech if *p* is 1, albeit with a harsh, noisy timbre). This procedure enabled us to adapt the intelligibility of our speech stimuli (i.e. *p*) to the performance of each individual participant, as explained in the following section.

### Experimental Design

The experimental design and statistical procedure (see below) for Experiment 2 was pre-registered (https://osf.io/peycm/; January 2018) prior to data collection (in February and March 2018). Data for Experiment 1 was collected prior to the pre-registration of Experiment 2 (in August and September 2017).

The experimental design was identical for Experiments 1 and 2, apart from the electrical stimulation protocol, described in the next section. Both experiments consisted of adaptive runs (A) in which task difficulty was adapted to individual performance, tACS runs (Stim), and sham stimulation (Sham) runs, explained in the following. The order of runs (Fig. 1C) was A-Stim-Sham-Sham-Stim-A-Sham-Stim-Stim-Sham, repeated twice. This design led to 4 adaptive runs, 8 Stim runs, and 8 Sham runs for each participant. Each run consisted of 32 trials. Each trial (Fig. 1A) consisted of the presentation of one noise-vocoded five-word sentence (described in Section Stimuli), 1.6 s long, followed by 5.44 s of silence during which participants indicated the words they had detected in the preceding sentence. This was done by using a mouse to click on the corresponding image on the screen (8-alternative forced choice for each of three word categories “number”, “color”, and “animal”; the first and last words were always “pick” and “up”, respectively, and did not need to be reported; see Section Stimuli and Fig. 1A). The experiment proceeded automatically if participants did not choose all required items in the allocated time interval.

In adaptive runs, the mixing proportion of 16- and 1-channel vocoded speech envelopes (*p;* defined in Section Stimuli) was adapted to the individual participant’s performance so that on average there was a probability of 0.5 that each of the three words in a given trial were identified correctly. This was achieved using the threshold estimation procedure that is part of the Psychtoolbox (Brainard, 1997) in MATLAB (The MathWorks), building on a method described by Watson and Pelli (1982). No stimulation was applied in these adaptive runs. The estimated mixing proportion was kept constant in the four (two Stim and two Sham) runs following the adaptive run and then updated in the next adaptive run, in order to account for any learning or fatigue effects. The outcome of our adaptation procedure, i.e. mixing proportions for individual participants and for each of the four adaptive runs, is shown in Fig. 1D.

In Stim runs, tACS was applied at 3.125 Hz such that neural oscillations should follow the alternating current (Herrmann et al., 2013). By generating tACS and acoustic waveforms from the same sound card we were able to control the phase relation between neural oscillations (reflected by the applied current) and speech rhythm. We assessed the effect of 8 phase relations (between 0 and 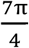, in steps of 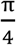; corresponding to delays between 0 and 280 ms, in steps of 40 ms for speech at 3.125Hz) between tACS and speech. This was done by applying tACS continuously and varying the timing of stimulus presentation so that the p-centers of the individual words were aligned with a certain tACS phase (dashed lines in Fig. 1A). In order to minimize physical sensations associated with the stimulation (Vosskuhl, Huster, & Herrmann, 2015), the current was ramped up and down for 3 s before and after each Stim run, respectively, following a Hanning window. Note that tACS is a silent technique; additional acoustic stimulation, which can entrain neural oscillations in addition to the applied current (e.g., Lakatos et al., 2008), can therefore be ruled out.

Sham stimulation was necessary to test (1) for generic effects of tACS (i.e. disrupting or enhancing, irrespective of tACS phase) and (2) whether a certain phase relation between tACS and speech enhances or disrupts speech perception (or both), relative to an unstimulated baseline (see Data Analyses). In Sham runs, the current was ramped up and down immediately (6 s in total) at the start of each test run. This had the goal of creating the usual sensations associated with tACS, but without stimulation during the remainder of the test run. The speech sequences were presented with the same delays as in the Stim runs that were necessary to vary the phase relation between tACS and speech rhythm in the latter condition.

In both Stim and Sham runs, the number, color and animal words presented in different trials were counterbalanced across the 8 phase relations between tACS and speech rhythm. This ensured that all words occurred equally often in all phase bins (and equivalent pseudo-phase bins for Sham runs). In total, 32 trials were tested in each phase bin such that 96 word report measures were obtained for each phase bin in each condition (Stim and Sham).

### Electrical Stimulation

Current was administered using one or two (depending on the experiment, see below) battery-driven stimulators (DC-Stimulator MR, Neuroconn GmbH, Ilmenau, Germany). Each of the stimulators was driven remotely by the output of one channel of a high-quality sound card (Fireface, UCX, RME, Germany); another output channel was used to transmit diotic auditory stimuli to the participants’ headphones (ER2 Earphones, Etymotic Research Inc., USA), assuring synchronization between applied current and presented stimuli.

We tested two different stimulation setups in the two experiments (Fig. 1B). In Experiment 1, we used the setup previously described in Zoefel, Archer-Boyd, et al. (2018). In this way, we were able to test behavioural consequences of a stimulation protocol for which neural effects have been described (see Introduction). This setup used unilateral (left hemisphere) stimulation, with two electrodes attached at location T7 and C3 of the 10-10 system, respectively. The size of the electrode over T7 (30 x 30 mm) was reduced as compared to that over C3 (50 x 70 mm). This had the goal of increasing the relative impact on oscillatory entrainment beneath the smaller more ventral electrode which was intended to be placed directly over auditory brain regions. Current intensity was set to 1.7 mA (peak-to-peak).

In Experiment 2, we used bilateral stimulation and replaced the standard rectangular electrodes with ring electrodes which have been shown to improve the focality and efficacy of the stimulation (Datta et al., 2009; Heise et al., 2016; Saturnino et al., 2015). Each pair of ring electrodes consisted of an inner, circular, electrode with a diameter of 20 mm and a thickness of 1 mm, and an outer, “doughnut-shaped”, electrode with an outer and inner diameter of 100 and 75 mm, respectively, and a thickness of 2 mm. The inner electrodes were centred on T7 and T8 of the 10-10 system, respectively. The parts of the outer electrodes which overlapped with participants’ ears were covered using isolating tape (Fig. 1B). As the total electrode surface (per hemisphere) was reduced to approximately 300 mm^2^ as compared 440 mm^2^ in Experiment 1, we also reduced current intensity to 1.2 mA (peak-to-peak), in order to avoid an increased likelihood of sensations associated with the stimulation. For both setups, electrodes were kept in place with adhesive, conductive ten20 paste (Weaver and Company, Aurora, CO, USA).

After each Stim or Sham run, participants were asked to rate the subjective strength of any sensations induced by the stimulation by giving a 10-point rating between 0 (no subjective sensations) and 10 (strong subjective sensations). Although sensation ratings were relatively low in all conditions and experiments, Stim runs (Experiment 1: 1.80 ± 1.45; Experiment 2: ± 1.62; mean ± SD) were rated significantly higher than Sham runs (Experiment 1: 1.01 ± 1.08; Experiment 2: 1.23 ± 1.27) in both experiments (Experiment 1: t(26) = 5.33, p < 0.001; Experiment 2: t(18) = 6.20, p < 0.001; paired t-test). However, even though participants were able to distinguish stimulation from Sham runs (cf. Turi et al., 2019), this is extremely unlikely to have influenced the critical behavioural outcome of stimulation. Participants would also need to distinguish different tACS phases, and relate these to the rhythm of speech signals, in order for there to be any influence of tACS phase on word report accuracy (cf. Discussion).

### Data Analyses

Data and analysis scripts are available in the following repository: https://doi.org/10.17863/CAM.43617

We applied two different statistical analyses to data for both experiments: A pre-registered analysis, and an analysis optimized based on simulations (Zoefel, Davis, Valente, & Riecke, 2019), both described in the following. Although data for Experiment 1 was collected prior to the pre-registration of Experiment 2, the pre-registered analysis had not been applied to the data at that time. The optimized analysis was selected exclusively based on results from simulations (Zoefel et al., 2019) and had not been applied to any of the datasets prior to selection.

The following analysis protocol was pre-registered for Experiment 2:

1. Performance in the word report task (proportion correct) was calculated, separately for each target word. In the Stim condition, this was done separately for the 8 phase relations between tACS and speech rhythm. In the Sham condition, word report accuracy was averaged across the 8 (pseudo-) phase relations, as word report accuracy cannot depend on tACS phase (since no tACS was applied).
2. Despite our use of adaptive runs to ensure that word report probability (averaged across target words) was 0.5, some words were more accurately reported than others. This was to be expected based on previous findings showing that some speech sounds (e.g. fricatives) and some phonetic features (e.g. voicing) are more readily perceived in vocoded speech (Shannon et al, 1995). Our analysis protocol therefore needed to take this baseline variation in accuracy into account (since a tACS-induced improvement in word report accuracy from, e.g., 90% to 95% reflects a larger change in perception than an improvement from, e.g., 50% to 55%). To normalize performance changes for each target word by baseline accuracy, we therefore quantified the magnitude of tACS effects by calculating the Log Odd’s Ratio (LOR) between Stim and Sham conditions:

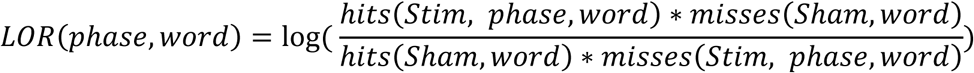 In this analysis, *hits* and *misses* are the number of correctly and incorrectly identified words, respectively, in each single condition (4 trials per phase and word in the Stim condition, 32 trials per word in the Sham condition since phase is irrelevant). LOR is 0 if there is no difference between Stim and Sham conditions (i.e. no effect of tACS on word report), and negative or positive if tACS disrupts or enhances word report accuracy (for a given tACS phase and word), respectively. LOR was then averaged across all of the 24 target words. The null hypothesis states that LOR is not modulated by the phase relation between tACS and speech rhythm. Target words were excluded from the analysis if they were never identified in any of the (Stim and Sham) conditions, as such a low word report probability might have prevented any consequence of tACS, even if effective. If a target word was never identified for a given tACS phase in the Stim condition, but was identified at least once in the corresponding Sham condition, then 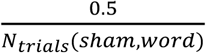(i.e.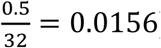) was added to *hits(Stim, phase, word)* and *misses(Stim, phase, word)*, and 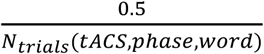 (i.e. 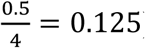 was added to *hits(Sham, word)* and *misses(Sham, word)*. This is a modified version of a procedure proposed by, e.g., Macmillan & Creelman (2004) and ensures that the change in accuracy level (i.e. LOR) in the Stim condition (i.e. the added 0.0156 hits or misses relative to the number of trials: 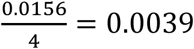) is the same as that in the Sham condition 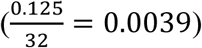.
3. The “preferred” phase relation between tACS and speech rhythm was defined as the maximal LOR. As expected from previous studies (e.g., Riecke et al., 2018; Zoefel, Archer-Boyd, et al., 2018), this varied across participants in a uniform fashion (Experiment 1: p = 0.34; Experiment 2: p = 0.85; Rayleigh’s Test for non-uniformity of circular data). Before the tACS-induced modulation of word report accuracy could be averaged across participants and quantified statistically, performance was therefore aligned at a common phase bin for each participant. We applied two alignment procedures prior to statistical analysis (Fig. 3), as described in the following.

**Figure 3.**
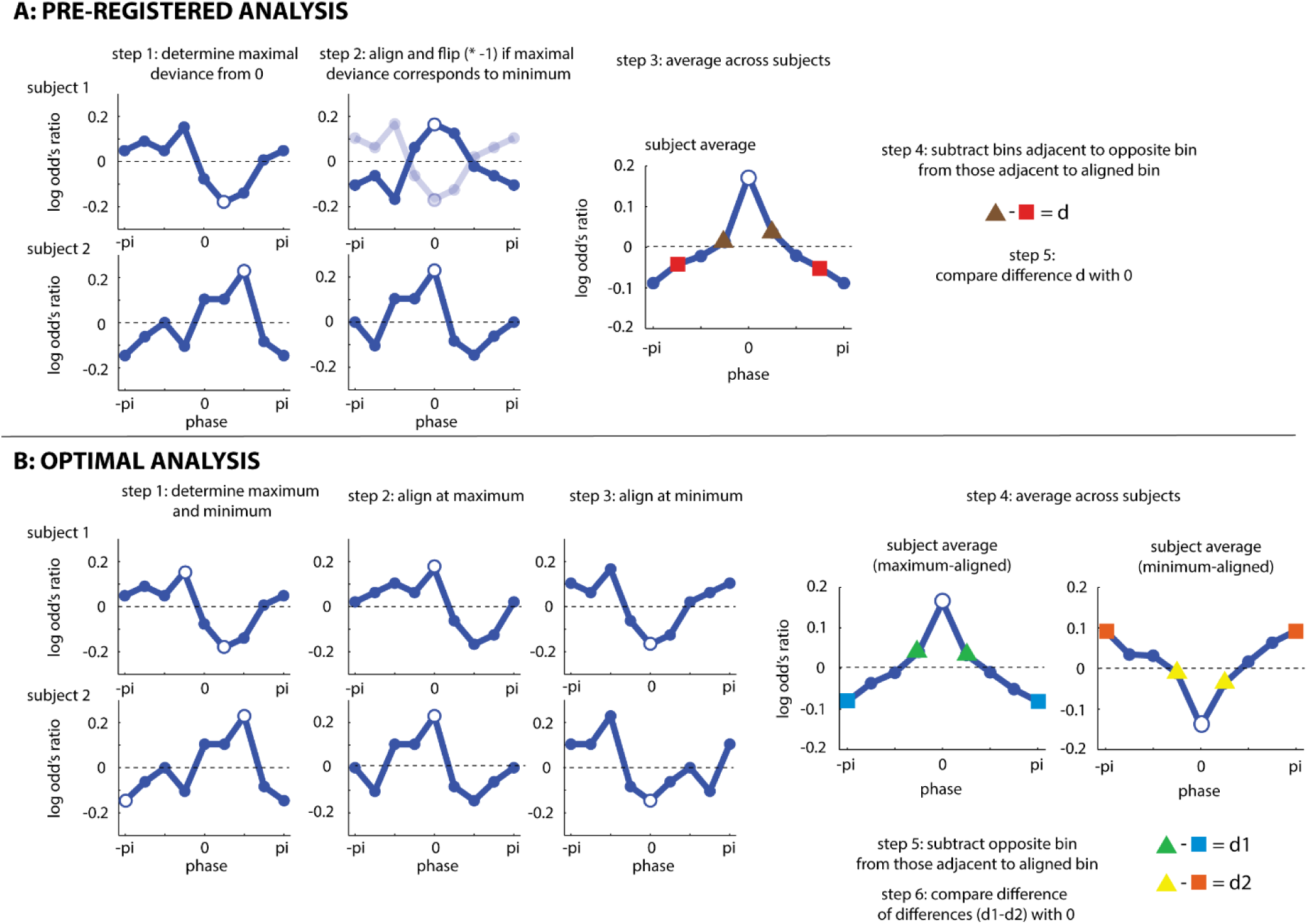
Statistical Protocols. Both analyses (**A**: pre-registered analysis; **B**: optimal analysis, identified in Zoefel et al., 2019) are illustrated based on two (simulated) example subjects, and the average across 20 (simulated) subjects. Open circles show phase bins used for alignment. In all panels, the pi/-pi bin is plotted twice for visualization purposes.

#### Pre-registered procedure for detecting effects of tACS phase on word report (Fig. 3A)

Performance (the LOR between Stim and Sham conditions) was quantified as a function of the phase relation between tACS and speech rhythm, as described above. Most previous studies average data across participants by aligning maximal performance at a certain phase bin (e.g., the center bin) and subsequently phase-wrapping the remaining data for each participant. However, given previous results concerning the suppression of the BOLD response by tACS (Zoefel, Archer-Boyd, et al., 2018), it is equally plausible that tACS disrupts rather than enhances word report accuracy; in this case, it might be more appropriate for the *minimum* performance to be aligned over participants. Thus, our pre-registered approach was that for each participant, we would designate as the center bin the phase bin that most strongly deviated from a LOR of 0 (i.e. the maximum *or* minimum, whichever had the highest absolute value). If aligned performance corresponded to a minimum for a certain participant, the sign of the resulting LOR values was flipped (i.e. multiplied by −1) so that the value aligned at the center bin was consistently a maximum across participants. It is possible that tACS also modulates speech perception in a phase-independent manner (i.e. enhances or disrupts performance for *all* phase bins). This generic effect of tACS would result in an average LOR (combined over bins) that is larger or smaller than 0 (respectively). We removed this effect from individual participant’s data by subtracting the mean LOR over phase bins. This mean correction operation was performed before the phase bin with the largest deviance from 0 was determined. This final step was not pre-registered but deemed necessary post-hoc, as the largest deviance from 0 would otherwise always reflect the direction of the generic effect (i.e. it would always be a maximum given a positive generic effect, and always a minimum given a negative generic effect).

Due to the alignment procedure described above, performance in the center bin is trivially a maximum and needs to be excluded from the analysis. However, assuming that tACS modulates speech perception in a phase-specific manner, performance in phase bins next to the center bin should still be better than in phase bins that are further away. In order to test whether word report accuracy depends on the phase relation between tACS and speech, we therefore subtracted the average LOR of the two phase bins adjacent to the bin that is 180° away from the center bin (red in Fig. 3A) from that of the two phase bins adjacent to the center bin (brown in Fig. 3A). The two phase bins adjacent to the bin opposite to the center bin, rather than the opposite bin itself, were analyzed so that our comparison involved an equal number of phase bins. Separately for each of the two experiments, the resulting difference *d* was then compared to 0, using a one-tailed one-sample t-test. A *d* greater than 0 represents evidence for a causal role of neural entrainment for speech perception. This is a variant of an analysis used by Riecke et al. (2018, Experiment 1). To test whether phasic modulation of speech perception depends on electrode configuration (unilateral vs bilateral), *d* was also compared across the two experiments, using a two-tailed two-sample t-test. This comparison across experiments was not pre-registered.

#### Optimized procedure for detecting effects of tACS phase on word report (Fig. 3B)

Even though our pre-registered procedure is statistically valid (i.e. not prone to false-positives; Asamoah, Khatoun, & Mc Laughlin, 2019a), and was similar to analyses that had been used in published papers at the time of pre-registration, it was unclear whether this was the optimal approach for detecting tACS-induced phase effects. Subsequently, and in parallel with data collection, we ran Monte-Carlo simulations designed to determine the optimal analysis for this and similar scientific questions (Zoefel et al., 2019). In this study, we simulated 2 x 1000 datasets with and without a phase effect on binary perceptual outcomes (“hit” vs “miss”). We then applied different analyses and tested which of these was best able to detect true effects, and reject absent phase effects on perceptual report. The optimal analysis was determined for combinations of different parameters that either concern the nature of the underlying phase effect on perception (effect size, effect width, effect asymmetry) or the experimental paradigm used to measure the phase effect (number of trials, number of phase bins tested etc.). Based on these simulation results, we were able to determine the analysis optimally suited to analyze the data collected in the current study. Importantly, and unfortunately, the optimal analysis was not the one that we pre-registered. Indeed, our pre-registered analysis was never the optimal approach for any of the combinations of parameters tested.

The optimal analysis (MAX-OPP VS MIN-OPP) described below for analyzing the present experiment was consistently preferred for: (1) findings averaged across all tested effect widths and asymmetries, (2) weak effect sizes (4%-8% peak-to-peak modulation of performance, cf. Riecke & Zoefel, 2018), (3) dichotomous outcome measures (“hit” vs “miss”), and (4) the experimental parameters used in our study (8 tested phases and 256 trials with 3 target words reported in each trial). Note that, although simulations showed methods based on regression with circular predictors and/or permutation tests to be superior in general, these could not be used in the present study. First, target words in the present study were counterbalanced across tACS phase bins; this was important to control for item-specific variation in report accuracy described above. Random assignment of trials to phase bins, as required for permutation tests, would destroy this counterbalanced design and potentially lead to invalid results when comparing observed and null phase effects. Second, again due to item-specific variation, we quantified tACS effects using LOR, i.e. expressed as word report accuracy normalized by baseline accuracy for each target word (see above). As LOR is not defined on the single-trial level, we were unable to apply methods regressing single-trial responses on circular predictors.

Based on these selection criteria, we determined that the MAX-OPP VS MIN-OPP analysis in Zoefel et al. (2019) was optimal and we therefore applied this analysis to the data from the current experiment.

The maximal performance of individual participants was aligned at the center bin, and the remaining bins phase-wrapped. Performance in the bin opposite (i.e. 180’ away from) the center bin (blue in Fig. 3B) was subtracted from the average of the two bins adjacent to the center bin (green in Fig. 3B), yielding difference *d1*. If a tACS phase effect were present, we would expect a relatively high value for *d1*. The procedure was then repeated, but this time using data aligned to minimal performance. Again, performance in the bin opposite to the center bin (orange in Fig. 3B) was subtracted from the average of the two bins adjacent to the center bin (yellow in Fig. 3B), yielding difference *d2*. If a tACS phase effect were present, we would expect a negative value for *d2*. Separately for each of the two experiments, the difference between *d1* and *d2* was then compared against 0, using a one-tailed one-sample t-test. A difference greater than 0 represents evidence for a causal role of neural entrainment for speech perception. To test whether phasic modulation of speech perception depends on electrode configurations (unilateral vs bilateral), the obtained difference was also compared across the two experiments, using a two-tailed two-sample t-test.

#### Enhancing vs disrupting word report

The analyses described above detect phase-specific effects of tACS on word report and therefore provide evidence that tACS-induced changes in neural entrainment modulate speech perception. However, they cannot answer the question of whether the effects observed reflect *enhancement* or *disruption* of speech perception (or both) relative to the Sham condition. Additional analyses were therefore carried for this purpose, as described in the following.

#### Pre-registered procedure

For the pre-registered analysis, the maximum of individual LOR data was aligned at the center bin and remaining bins were phase-wrapped. Separately for each of the two experiments, the average of the two bins (green in Fig. 3B) adjacent to the center bin (which is trivially a maximum and cannot be analyzed) was compared to 0, using a one-tailed, one-sample t-test. A LOR greater than 0 reflects an enhancing effect of tACS on word report relative to Sham. This average was then compared across experiments, using a two-tailed, two-sample t-test, to test whether the hypothetically enhancing effect depends on electrode configuration (unilateral vs bilateral). This procedure was repeated, but the minimum LOR was aligned at the center bin before the average of the two adjacent bins (yellow in Fig. 3B) was compared against 0. A LOR smaller than 0 reflects a disruptive effect of tACS on word report relative to Sham. This average was again compared across experiments, to test whether the hypothetically disruptive effect depends on electrode configuration. This comparison across experiments was not pre-registered.

Importantly, this analysis procedure tests for extreme performance values (maxima and minima). However, the actual extreme values are used for the alignment of individual data to a common phase bin and therefore need to be excluded from subsequent analysis. This pre-registered approach might therefore increase the probability of a false negative result: Enhancing or disrupting effects might exist but cannot be revealed as only the second most extreme values are compared against 0. We therefore also report a potentially more optimal procedure, inspired by the outcome of our Monte-Carlo simulations (Zoefel et al., 2019).

#### Optimized procedure

Data was aligned using maximal or minimal performance, as described. However, instead of comparing the average of the two bins adjacent to the center bin against 0, we compared the opposite bin against 0 (using one-tailed, one-sample t-tests). If tACS-induced changes in entrainment facilitated word report, then this opposite bin (orange in Fig. 3B) should be larger than 0 after alignment to the phase bin with the minimum LOR. If tACS-induced changes in entrainment disrupted word report, then this opposite bin (blue in Fig. 3B) should be smaller than 0 after alignment to the phase bin with the maximum LOR. In addition to the comparison against 0 for each of the experiments separately, respective data was again compared across experiments (using two-tailed, two-sample t-tests), to test whether the observed enhancement/disruption depends on electrode configuration (unilateral vs bilateral).

#### Bayesian Statistics

For most statistical tests, we also report the Bayes Factor for the presence (*BF*_10_) or absence *BF*_01_; reported if BF_10_ < 1) of an effect, respectively, where 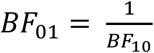. The bayesFactor toolbox for MATLAB was used for this purpose (https://klabhub.github.io/bayesFactor/).

## Results

We used two different statistical analyses to test whether tACS-induced changes in neural entrainment causally modulate the perception of isolated words in rhythmic sentences. The first analysis (Fig. 3A) was pre-registered prior to data collection for Experiment 1, but was found to be sub-optimal in a simulation study design to reveal optimal analyses for oscillatory phase effects (Zoefel et al., 2019). The second analysis (Fig. 3B) was not pre-registered but was identified as the optimal analysis for our purposes based on the latter study (see Section Data Analyses).

### Modulation of word report accuracy by tACS-induced changes in neural entrainment

In both experiments, the probability of correctly identifying each of the three target words (averaged across tACS phases) was close to 0.5 (Experiment 1: Stim, 0.51 ± 0.07, Sham, 0.51 ± 0.07; Experiment 2: Stim, 0.47 ± 0.13; Sham, 0.46 ± 0.13; mean ± SD), and not different between stimulation and sham conditions (Experiment 1: t(26) = 0.41, p = 0.68; Experiment 2: t(18) = 0.61, p = 0.55; paired t-test). This shows that our procedure to adjust individual word report accuracy to near-threshold level was successful (see Section Experimental Procedure), and that there is no net effect of tACS when word report data is combined over different phase relations. The following analyses are based on the difference in word report accuracy between stimulation and sham conditions as a function of phase relation, quantified using Log Odd’s Ratio (LOR; see Section Data Analyses).

Fig. 4A-C shows LOR as a function of the phase relation between tACS and speech rhythm. Data was aligned in three different ways before being averaged across participants, as required for the different statistical analyses (see Section Data Analyses), and is shown separately for the two stimulation protocols (unilateral in Experiment 1 vs bilateral in Experiment 2). Panels shaded grey in Fig. 4 are relevant for the pre-registered analysis; the other panels concern the optimized analysis.

**Figure 4.**
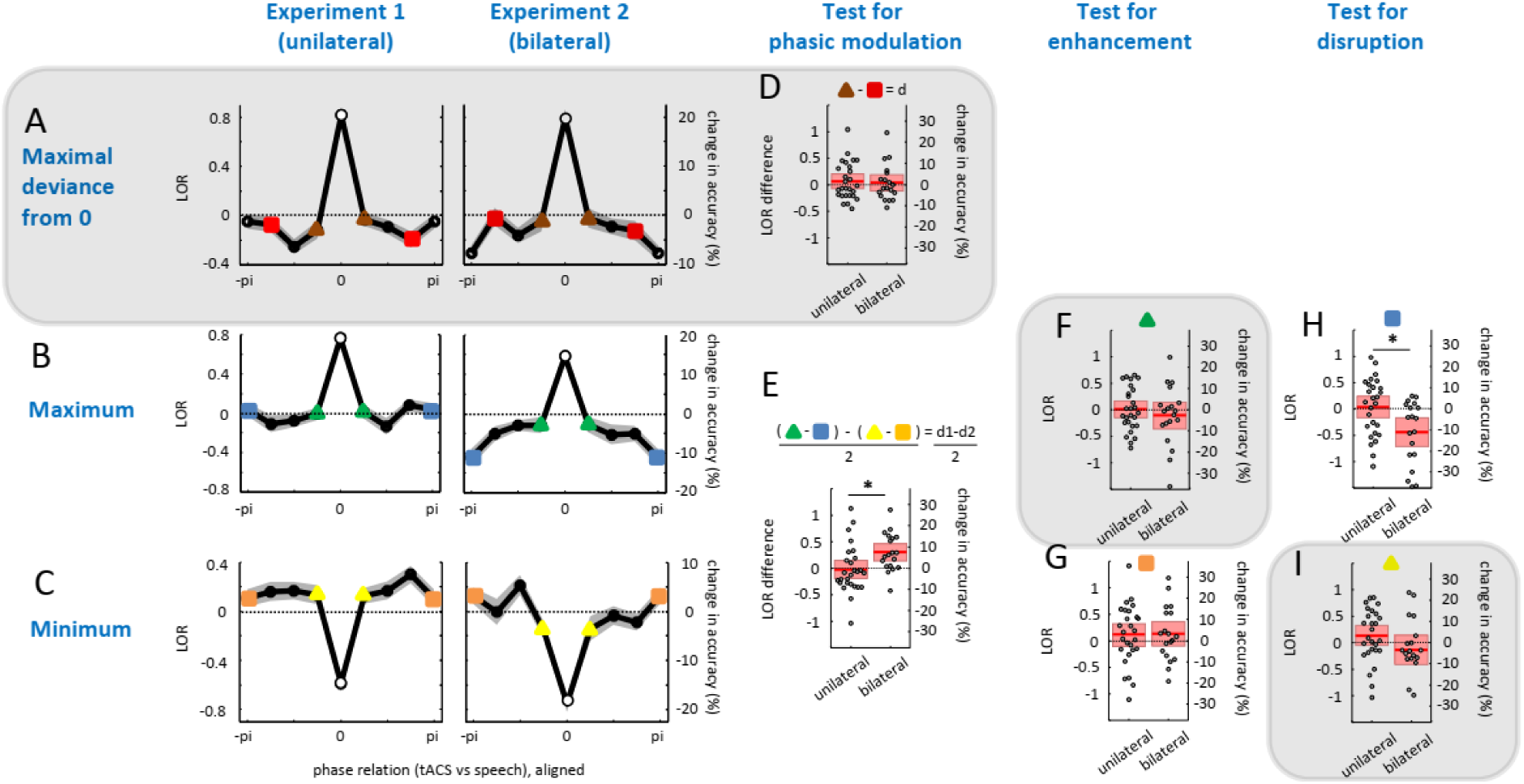
Main results. **A-C.** Average word report accuracy (as Log Odd’s Ratio, LOR) as a function of the phase relation between tACS and speech rhythm. As required by the two applied statistical analyses (see Section Data Analyses and Fig. 3), individual data was aligned in different ways before being averaged across participants, shown in rows A-C. Bins used for alignment, by definition maximal or minimal, are shown as open circles. Shaded areas show standard error of mean (SEM) after between-participant variation has been removed as appropriate for repeated-measures comparisons of different phase bins (Cousineau, 2005). The pi/-pi bin is plotted twice for visualization purposes. **D-I**. Distribution of values (relevant phase bins are color-coded in A-C) which are compared to 0 to test for a phasic modulation (D,E), enhancement (F,G), or disruption (H,I) of speech perception, respectively. Dots show data from single participants, mean and confidence interval (1.96*SEM) are shown by red lines and red-colored areas, respectively. In all panels, right y-axes show LOR converted into changes in word report accuracy, given, on average, 50% correctly identified target words. Note that these changes are expressed relative to the Sham condition (A-C, F-I) or relate two phase bins in the Stim condition (D,E). In panel E, LOR difference values (d1-d2) and corresponding changes in accuracy were divided by two, to take into account the fact that this difference involves two comparisons of phase bins (d1, cf. panel B, and d2, cf. panel C). This was not necessary for the corresponding statistical test (see Section Data Analyses) which is unaffected by such scaling factors. Figure panels shaded grey correspond to those relevant for the pre-registered analysis.

When using our pre-registered analysis (Fig. 4D), we did not find strong evidence for a modulation of word report by the phase relation between tACS and speech rhythm, neither for unilateral (t(26) = 0.89, p = 0.19; *BF*_01_ = 2.11; the Bayes factor notation indicates which hypothesis is supported, see Methods) nor bilateral stimulation (t(18) = 0.42, p = 0.34; *BF*_01_ = 2.93). There was no reliable difference between stimulation protocols (t(44) = 0.26, p = 0.80; *BF*_01_ = 3.29). When using the analysis that was identified to be optimal using simulations (Fig. 4E; Zoefel et al., 2019), we found very strong evidence for a tACS-induced modulation of speech perception for bilateral stimulation (t(18) = 3.65, p = 0.0009; *BF*_10_ = 44.19). There was evidence for the absence of an effect for unilateral stimulation (t(26) = −0.30, p = 0.62; *BF*_01_ = 6.15). The observed modulation of speech perception in the bilateral stimulation condition was stronger than that in the unilateral condition (t(44) = 2.60, p = 0.01; *BF*_10_ = 4.10). The right y-axes in Fig. 4 show how LOR relates to changes in word report accuracy, given an average word report of 50 %. We estimate that a change from a near-optimal phase relation between bilateral tACS and speech (green and yellow phase bins in Fig. 4) to a non-optimal phase relation (blue and orange bins) leads to an ∼8% change in word report accuracy (Fig. 4E).

### Enhancing vs disrupting word report

We also used two different analyses to determine whether tACS-induced changes in neural entrainment enhance or disrupt speech perception (or both) relative to the Sham condition: A pre-registered analysis and an analysis optimized based on theoretical considerations (described in Section Data Analyses).

For most conditions, our pre-registered analysis revealed evidence for an absence of enhancing (Fig. 4F) or disruptive (Fig. 4I) changes in word report induced by the phase relation between tACS and speech rhythm, for both unilateral (enhancing: t(26) = 0.04, p = 0.48; *BF*_01_ = 4.77; disruptive: t(26) = 1.33, p = 0.90; *BF*_01_ = 11.43) and bilateral stimulation (enhancing: t(18) = 0.88, p = 0.80; *BF*_01_ = 7.64; disruptive: t(18) = 1.01, p = 0.16; *BF*_01_ = 1.60). There was no reliable evidence that the two stimulation protocols differ in their enhancing (t(44) = 0.81, p = 0.42; *BF*_01_ = 2.60) or disrupting (t(44) = 1.65, p = 0.11; *BF*_01_ = 1.15) effect.

Using our optimized analysis, the unilateral condition showed some evidence for an absence of enhancement (Fig. 4G) and disruption (Fig. 4H) of speech perception (enhancing: t(26) = 0.99, p = 0.17; *BF*_01_ = 1.89; disruptive: t(26) = 0.32, p = 0.63; *BF*_01_ = 6.24). Strong evidence for disruption of word report performance was observed in the experiment using bilateral stimulation (t(18) = −3.23, p = 0.002; *BF*_10_ = 19.67), whereas it remained inconclusive whether speech perception can be enhanced (t(18) = 1.07, p = 0.15; *BF*_01_ = 1.50). Disruption of speech perception caused by bilateral stimulation was stronger than that in the unilateral condition (t(44) = 2.81, p = 0.007; *BF*_10_ = 6.24), but the unilateral and bilateral experiments did not reliably differ in their enhancing effect on word report (t(44) = 0.12, p = 0.91; *BF*_01_ = 3.36). In the bilateral experiment, the mean LOR for the phase bin used to test for disruptive effects (Fig. 4I) was −0.44 (± 0.60). Given an average word report of 50%, this translates into an average reduction of ∼11 % accuracy at this particular tACS phase, relative to the sham condition.

## Discussion

The present study replicates previous findings that the timing of tACS, synchronized with the amplitude envelope of spoken sentences, can change speech processing (Riecke et al., 2018; Wilsch et al., 2018; Zoefel, Archer-Boyd, et al., 2018). In addition, several aspects of the study are novel and go beyond the current state-of-the-art.

First, it was previously unclear whether observed effects reflect tACS-induced changes in listeners’ ability to segregate speech and background noise (i.e. auditory stream segregation) or a modulation of processes that are more central to speech perception (such as the perception of speech sounds and recognition of words). Modulation of any of these processes could lead to the tACS-induced changes in word report shown previously by Riecke et al. (2018) and Wilsch et al. (2018). In this study, we demonstrate that the phase relation between tACS and rhythmic speech affects word report accuracy, even for sequences of degraded words presented in silence. Although noise-vocoded speech, used in this study sounds “noisy”, it is a form of intrinsically degraded speech. Consequently, there is only one sound source present and no other external signal (such as background noise) must be segregated to achieve effective perception (see Mattys et al., 2012, for further discussion of the distinction between intrinsic and extrinsic degradation of speech). The present findings go beyond pre-existing results by showing that neural entrainment can causally modulate central processes that support speech perception without any requirement for auditory scene analysis (e.g., segregation of competing speakers or background noise).

Second, previous work (Experiment 2 in Riecke et al., 2018; Wilsch et al., 2018) used natural speech, which has a dominant perceptual rhythm (see Fig. 1C in Wilsch et al., 2018) that is conveyed by a complex quasi-periodic amplitude envelope with considerable variation in the modulation spectrum (Chandrasekaran, Trubanova, Stillittano, Caplier, & Ghazanfar, 2009; Peelle & Davis, 2012; but see Cummins, 2012). In these previous studies, perception was modulated when tACS waveforms were delayed relative to the speech signal, with best word report at a delay of 375 ms in Riecke et al. (2018), and 100 ms in Wilsch et al. (2018). These inconsistent best time delays might be a consequence of the use of quasi-periodic natural speech, which leads to inconstant or unspecified phase shifts between tACS and speech. In contrast, the use of perfectly rhythmic speech (in perceptual terms) allowed us to specify the phase lag between speech and brain stimulation more precisely. In this work, we define zero phase as occurring at the perceptual rhythm of speech (i.e. at the p-centers of each syllable, see Methods). This illustrates a method by which links between the amplitude envelope for speech, neural oscillations and perceptual rhythms can be tested (cf. Oganian & Chang, 2018).

A third and final area of novelty is that we compared different analysis methods and electrode configurations for tACS studies of speech perception. Our findings from these methodological explorations have important implications for future work which we will discuss in detail in the next section.

### Methodological Implications

Demonstrating a causal role of neural entrainment for speech processing is an important step forward from previous correlational studies (Gross et al., 2013; Peelle et al., 2013; Zoefel & VanRullen, 2015a) which show a link between intelligibility and entrainment but are unable to establish the causal direction between these factors. While positive results in published studies provide evidence of causality, it remains important to optimize our methods if we to demonstrate causality beyond reasonable doubt: Otherwise any subsequent failure to replicate might be mistakenly interpreted as an absence of a causal role of these mechanisms on underlying neural processes (e.g., speech perception). Furthermore, concerns have already been raised concerning analytic methods used in existing studies (e.g., Asamoah et al., 2019a), and the possibility of peripheral effects of electrical stimulation (e.g., Asamoah, Khatoun, & Mc Laughlin, 2019b).

tACS is one of only a few non-invasive techniques which can be used to manipulate oscillatory neural activity and/or entrainment in humans. Nevertheless, for obvious ethical and safety reasons (Fertonani, Ferrari, & Miniussi, 2015), the intensity of stimulation cannot be increased arbitrarily. Consequently, the current that actually reaches neural tissue is relatively weak (Huang et al., 2017; Opitz et al., 2016). This issue has led to ongoing debates about whether and how tACS can manipulate neural activity (Krause, Vieira, Csorba, Pilly, & Pack, 2019; Lafon et al., 2017; Opitz, Falchier, Linn, Milham, & Schroeder, 2017; Ruhnau, Rufener, Heinze, & Zaehle, 2018; Vöröslakos et al., 2018; Vosskuhl, Strüber, & Herrmann, 2018). Our study addresses two experimental variables – electrode montage and statistical analysis – which need to be carefully considered in tACS studies of speech processing.

First, it has often been suggested that the configuration of stimulation electrodes is crucial for the outcome of transcranial electrical stimulation experiments (Saturnino et al., 2015; Saturnino et al., 2017; Zoefel & Davis, 2017). However, very few studies have compared two different stimulation protocols for the same perceptual task. We found that word report accuracy was only modulated by the phase relation between tACS and speech rhythm if the stimulation was applied bilaterally using ring electrodes (but not unilaterally using square electrodes over the left hemisphere). Furthermore, we obtained a significant difference in between-experiment comparisons and can therefore conclude that the perceptual impact of tACS reliably differs between our bilateral ring and unilateral square electrode stimulation. As all other aspects of the experimental procedure were identical between the two experiments, these changes to the stimulation protocol are clearly relevant to neural entrainment and determine changes in perception. However, it is not clear whether the critical difference was the change in electrode shape, or bilateral stimulation. Although further investigations will be needed to address this point, we note that previous studies by Wilsch et al. (2018) and Riecke et al. (2018) used square electrodes and bilateral stimulation. This suggests that a bilateral electrode configuration might be key to obtaining reliable behavioural effects of tACS. Interestingly, we were able to use a unilateral tACS protocol to induce changes in the BOLD fMRI response to speech (Zoefel, Archer-Boyd, et al., 2018). This difference between neural and behavioural outcomes might arise if local changes in neural activity (measured voxel-by-voxel by fMRI) only lead to measurable changes in behavior if stimulation effects reach a sufficient number or proportion of functionally relevant voxels (e.g., in auditory brain regions) that are responsible for perceptual outcomes. This might explain why bilateral stimulation protocol is more effective since it plausibly leads to more widespread changes in neural activity.

Second, the analysis used to assess behavioural modulation of word report scores is also critical. We obtained reliable evidence of phasic modulation of word report using a parametric analysis method that was shown to be optimal in a recent simulation study (Zoefel et al., 2019). However, our pre-registered analysis, which was chosen before conducting these simulations, failed to reveal any reliable effect. Both analyses are scientifically valid (i.e. control for false-positives at the expected rate; cf. Asamoah et al., 2019a), and yet only one of these two seems to be sufficiently sensitive to reveal oscillatory phase effects. The pre-registration of experimental and analytical procedures is an important step to reduce the probability of false positive results and “p-hacking” (Ledgerwood, 2018; Nosek, Ebersole, DeHaven, & Mellor, 2018). However, the present work illustrates the challenges of pre-registering the right analysis for complex experimental designs in which the optimal analysis has not been clearly established by prior research or through simulations. In the absence of careful simulation to establish optimal analyses, null effects for pre-registered analyses may be too often observed.

### Does tACS enhance or disrupt speech perception?

One important motivation for exploring tACS-induced modulation of speech perception is that this might help populations that struggle to understand speech effectively, such as hearing-impaired listeners or individuals with developmental or acquired language impairments. One long-term goal for this work will therefore be to use tACS to *enhance* perception and comprehension of speech in challenging situations (cf. Peelle, 2018). Until now, however, the direction of phase-specific tACS effects (whether they enhance and/or disrupt word report) has been difficult to determine (Riecke & Zoefel, 2018).

Wilsch et al. (2018) found an improvement of ∼0.4 dB of speech-reception threshold (i.e. listeners can achieve 50% word report with a noise that is 0.4 dB louder with tACS than without) when comparing performance at individually “preferred” phases with a sham condition. However, it remains unclear whether this result reflects a true enhancing effect of tACS or is a false positive due to the way that the data was analyzed: Phase bins with highest performance were selected for the analysis of tACS but equivalent selection was not performed for the sham conditions (see Asamoah et al., 2019a, for relevant simulations and discussion).

Riecke et al. (2018, Experiment 1) avoided this bias by aligning maximum performance to a common phase bin over participants, then excluding this bin from further analyses. To test for an enhancing effect, they compared performance in the two phase bins adjacent to the bin used for alignment with sham. However, this analysis did not reveal an enhancing effect of tACS. This approach corresponds to our pre-registered method, which replicates their finding. To test for a disruptive effect, Riecke and colleagues extracted performance from the two phase bins adjacent to the bin opposite to the one used for alignment, and compared it with sham. They reported a disruption of speech perception in those phase bins.

Importantly, although the approach taken by Riecke et al (2018) is immune to the analytic bias identified by Asamoah et al. (2019), the exclusion of the phase bin corresponding to maximal performance, as well as the bin opposite to it, makes it difficult to test for a tACS-induced enhancement or disruption of performance, respectively. Our optimized approach (see Section Data Analyses) avoids this issue by testing for enhancement and disruption using phase bins that are opposite to that used to align data over participants. Aligning to minimum performance and maximum performance allows testing of enhanced and disrupted perception, respectively. This approach depends on assuming that phasic modulation of speech perception is such that maximum enhancement and maximum disruption occur in opposite phase bins. This assumption could be correct if a single neural population underlies both enhancement and disruption of speech perception, though more complex interactions between different neural populations are also possible.

The results obtained in the present study seem to confirm those reported by Riecke et al. (2018) and suggest that changes in neural entrainment, induced by tACS, lead to impaired speech perception relative to the sham condition (although note that Bayesian statistics remained inconclusive on the question of whether bilateral stimulation can enhance speech perception relative to sham). At face value, this might suggest pessimism in applying tACS to impaired individuals, for whom disrupting speech perception will be of limited practical value. However, recall that we only tested healthy participants who (presumably) have intact neural mechanisms for optimally entraining to speech. If we assume that healthy listeners can achieve optimal phase relation between neural activity and speech during natural listening (i.e. in our sham condition), then this might explain the limited evidence for tACS-induced enhancement of word report in our study. Conversely, impaired neural entrainment has often been reported for hearing-impaired listeners and older individuals (Henry, Herrmann, Kunke, & Obleser, 2017; Petersen, Wöstmann, Obleser, & Lunner, 2017; Presacco, Simon, & Anderson, 2016). It might therefore be that tACS can be used to restore optimal entrainment in impaired listeners who would not achieve optimal entrainment during natural listening. This possibility motivates further testing of tACS-induced enhancement of speech perception in impaired individuals.

### Limitations and Future Directions

Recent studies reported that participants can distinguish sham from transcranial current stimulation (Greinacher, Buhôt, Möller, & Learmonth, in press; Turi et al., in press), even if the former entails several seconds of stimulation, which is then faded out. Although we confirmed this finding in both experiments, we do not believe that placebo effects caused by these sensations can explain our findings. It is only if the sensations induced by the stimulation closely followed the rhythm of tACS, that they would produce an apparently phasic modulation of speech perception. It is however very unlikely that participants can distinguish the sensations caused by different tACS phases, and relate these to the rhythm of speech signals. First, participants typically report sensations such as itching or tingling, but no rhythmic component (at least at common stimulation intensities, such as those applied here), and these sensations seem to diminish or disappear relatively quickly after stimulation onset (e.g., Antal et al., 2017; Kessler, Turkeltaub, Benson, & Hamilton, 2012). Second, participants do not seem to be able to reliably distinguish different tACS frequencies (Nakazono, Ogata, Kuroda, & Tobimatsu, 2016; Wittenberg, Morr, Schnitzler, & Lange, 2019), including those which produce stronger sensations (10 or 20 Hz) than the frequency applied in the current study (3.125 Hz). Kleinert, Szymanski, & Müller (2017) also reported that participants are unable to determine whether frontal and parietal regions were stimulated in- or out-of-phase from each other. Nevertheless, a recent study convincingly demonstrated that some tACS effects on the motor system can be explained by stimulation of peripheral nerves (Asamoah et al., 2019b). It is possible that this issue could also affect stimulation of the auditory system, which needs to be addressed in future work.

A second issue to be explored in future work concerns the putative role of neural oscillations in mediating effects of tACS. Strictly speaking, the term “entrainment” merely describes a phenomenon in which a process becomes synchronized to an external rhythm (Pikovsky, 2008). Any neural or perceptual process that is modulated by the timing of tACS can therefore be considered “entrained”. However, a common, often implicit, assumption in the field of “neural entrainment” is that neural oscillations – endogenous rhythmic fluctuations in the excitability of neuronal ensembles (Buzsáki & Draguhn, 2004) – are involved. Neural oscillations are assumed to synchronize to an external rhythmic input, such as tACS, and yet a definitive demonstration that oscillatory activity underlies entrainment effects is not straightforward. The presentation of a regular stimulus (including speech) will evoke a regular sequence of (e.g., neural) responses and yield phenomena such as the “envelope following response” (e.g., Purcell, John, Schneider, & Picton, 2004). Similarly, tACS effects could be produced by the applied current interfering with speech processing, with the amount of interference determined by tACS phase and hence the magnitude and direction of current flow at that time. Future experiments need to be designed to resolve this issue: For instance, a tACS-induced modulation of speech perception which continues after stimulation would provide stronger evidence for an involvement of true neural oscillations (Zoefel, 2018; cf. Hickok, Farahbod, & Saberi, 2015; Kösem et al., 2018). Similar evidence for an involvement of endogenous oscillatory activity in the field of “entrainment” has been recently summarized elsewhere (Haegens & Zion Golumbic, 2018; Zoefel, ten Oever, & Sack, 2018).

As tACS was presented simultaneously with rhythmic speech sequences, the acoustic and electrical inputs were in competition to entrain neural oscillations. Given the fact that a large part of the applied current is shunted by skin and skull (e.g., Neuling, Wagner, Wolters, Zaehle, & Herrmann, 2012), it is likely that the presented speech was the dominant external force to entrain oscillations. If tACS is applied at an optimal phase relation to the speech stimulus, then this might boost the speech-induced entrainment (assuming it is not already at a maximum, as described above). If tACS is applied at the opposite (i.e. non-optimal) phase relation, it might decrease the speech-induced entrainment, leading to the reduced word report accuracy observed here. Nevertheless, how exactly acoustic and electrical stimulation interact has not yet been established and remains an exciting topic for future studies.

We found that the phase relation leading to highest (or lowest) word report accuracy was not consistent across participants (see Methods). This is a common finding in the tACS literature on speech processing (Riecke et al., 2018; Wilsch et al., 2018; Zoefel, Archer-Boyd, et al., 2018), and in neurophysiological recordings of auditory and visual perception (e.g., EEG studies from Busch, Dubois, & VanRullen, 2009; Henry & Obleser, 2012; Ng, Schroeder, & Kayser, 2012; VanRullen & Macdonald, 2012). However, it is still unclear what combination of functional anatomy and temporal properties of perceptual processing can explain these individual differences. Further research combining methods with high anatomical (e.g. fMRI) and temporal precision (e.g. EEG/MEG) will be required if we are to predict preferred phases for individual participants in tACS or similar studies.

## Acknowledgements

The authors thank Prof Axel Thielscher for helpful advice with the electrode configurations, and Simon Strangeways for help with their illustration. This work was supported by a grant from the German Academic Exchange Service (DAAD), the European Union’s Horizon 2020 research and innovation programme under the Marie Sklodowska-Curie grant agreement number 743482, and the Medical Research Council UK (grant number SUAG/008/RG91365).

## Conflict of Interest

The authors declare no competing financial interests.

## Notes

#### Summary of Updates

Extended discussion; included some bayesian statistics.

